# An Inflammatory and Quiescent HSC Subpopulation Expands with Age in Humans

**DOI:** 10.1101/2025.09.01.673389

**Authors:** Ksenia R. Safina, Dylan A. Kotliar, Michelle Curtis, Jonathan D. Good, Chen Weng, Shawn David, Soumya Raychaudhuri, Antonia Kreso, Jennifer Trowbridge, Vijay G. Sankaran, Peter van Galen

## Abstract

Aging of the blood system impacts systemic health and can be traced to hematopoietic stem cells (HSCs). Despite multiple reports on human HSC aging, a unified map detailing their molecular age-related changes is lacking. We developed a consensus map of gene expression in HSCs by integrating seven single-cell datasets. This map revealed previously unappreciated heterogeneity within the HSC population. It also links inflammatory pathway activation (TNF/NFκB, AP-1) and quiescence within a single gene expression program. This program dominates an inflammatory HSC subpopulation that increases with age, highlighting a potential target for further experimental studies and anti-aging interventions.

## Background

The aging blood system has a decreasing capacity to mount effective immune responses, transport oxygen, and produce lymphocytes^1–3^. Many of these changes can be traced to HSCs, which exhibit age-associated clonal expansion and myeloid bias^4–7^. Human studies across age groups have revealed multiple alterations including epigenetic reprogramming, decreased polarity, and skewed differentiation^8–11^. However, there is little consensus on the molecular programs consistently altered with age, in part due to technical variation across datasets^12^. Moreover, while the human HSC compartment is often analyzed as a whole, heterogeneity can be observed using single-cell sequencing^13,14^. The molecular basis of this heterogeneity, and how it evolves with age, remains poorly defined. To establish a robust reference for HSC aging and define the extent to which HSC heterogeneity changes with age, we undertook an effort to integrate human HSC aging studies. Our analysis reveals a consensus program of inflammation and quiescence in a subpopulation of HSCs that becomes increasingly dominant with age.

## Results and Discussion

To robustly characterize age-associated changes in human HSCs, we combined single-cell/single-nucleus data across six studies and one unpublished dataset from 98 individuals and annotated 28,989 HSCs using the BoneMarrowMap atlas (**Fig. 1a, Suppl. Table 1**). An orthogonal approach of identifying and annotating clusters on the integrated object yielded 44,187 HSCs and included 87% of HSCs identified by BoneMarrowMap (**Suppl. Fig. 1**), suggesting that the BoneMarrowMap algorithm selects an accurate and stringent population of HSCs. All downstream analysis was performed on BoneMarrowMap-annotated HSCs.

First, we inferred differential gene expression (DE) between young and aged samples with at least 20 HSCs (young: 19-37 y.o., n=25; aged: 60-87 y.o., n=15). 68 and 29 genes were up-and down-regulated in aged samples, respectively (**Fig. 1b, Suppl. Table 2**). Notably, three members of the AP-1 complex, *JUN, JUNB,* and *FOSB*, implicated in stress-activated MAPK signaling and quiescence, were consistently upregulated across studies (**Suppl. Fig. 2**).^15–17^

Gene set enrichment analysis (GSEA) revealed two major biological phenomena distinguishing expression profiles of young and aged samples (**Fig. 1c, Suppl. Table 2**). First, several inflammatory-response pathways, including TNF, IFN-g, and AP-1 signaling, were upregulated in aged samples. Second, aged samples showed higher quiescence, in contrast to young samples showing more active proliferation. These findings are in agreement with previous reports showing increased circulating TNF and delayed cell cycle entry of aged HSCs^9,18,19^.

**Figure 1.**
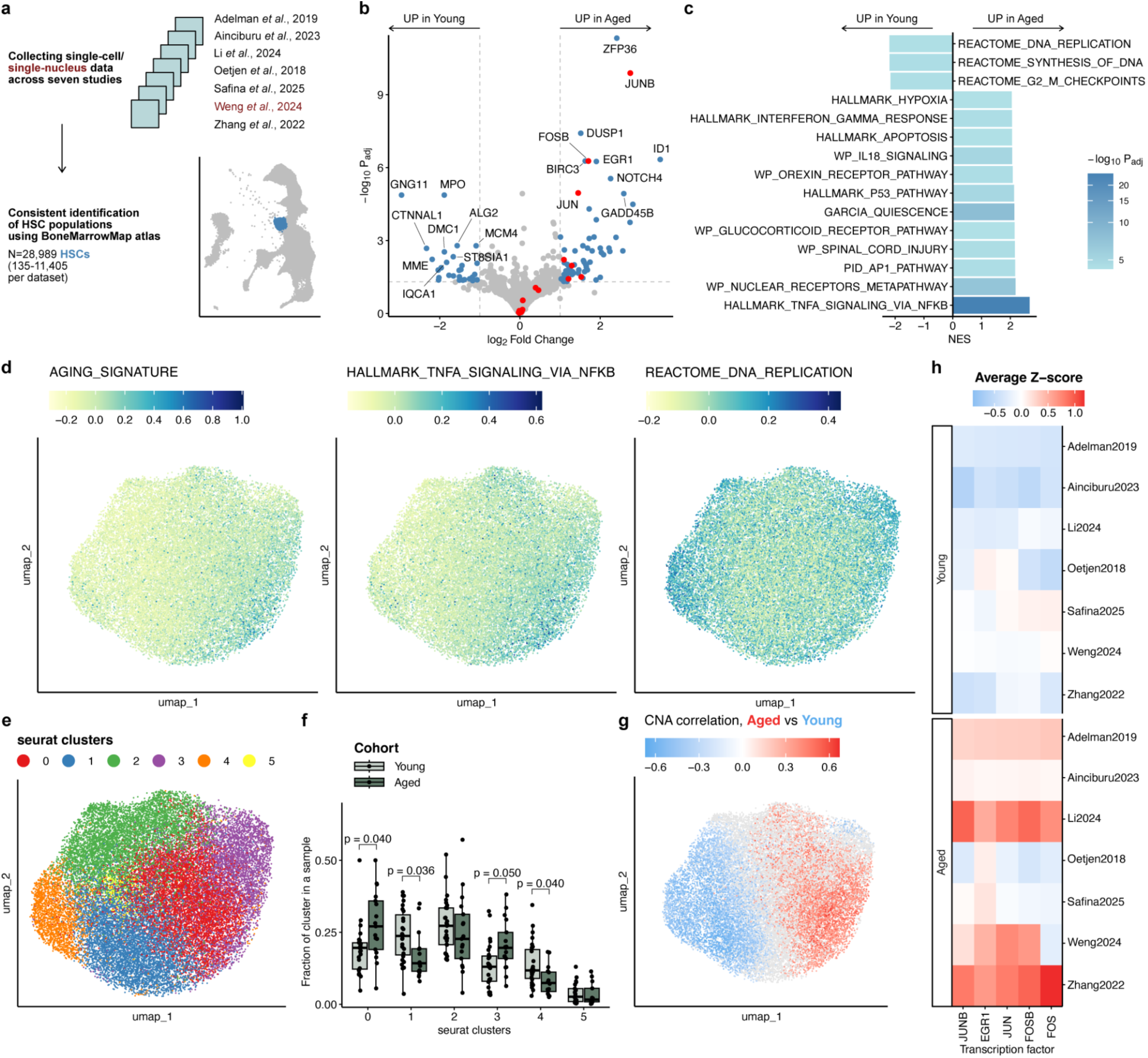
Data from multiple single-cell datasets show that transcriptional heterogeneity of human HSCs changes with age. **a.** Schematic shows dataset collection and HSC annotation. **b.** Volcano plot shows the differentially expressed genes in Young vs Aged samples across five datasets. AP-1 members are shown in red, genes with absolute logFC>1 and p.adjusted<0.05 are shown in blue; top-10 most significant hits per cohort and significant AP-1 members are labeled. **c.** Bar plot shows top-15 most significant gene sets in GSEA analysis; NES, normalized enrichment score. **d.** Uniform Manifold Approximation and Projections (UMAPs) show all integrated HSCs, colored by score for aging signature, Hallmark TNF-α/NF-κB signaling gene set, and Reactome DNA replication gene set. **e.** UMAP shows HSC clusters identified in Seurat. **f.** Box plot shows comparison of cluster abundances between the Young (n=27) and Aged (n=17) cohorts. Symbols indicate cluster fractions per sample; p-values of two-sided Wilcoxon test with Bonferroni correction are shown. **g.** UMAP is colored by neighborhood coefficient of correlation to Aged vs. Young cohorts; neighborhoods that didn’t pass FDR<10% are colored in gray. **h.** Heatmap shows activity of the top-5 regulons associated with the Aging cohort, identified by SCENIC; averaged Z-scores of AUC values are shown.

The activity of these pathways was not uniform across the population of HSCs (**Fig. 1d**, **Suppl. Fig. 3**). The aging signature (defined as 68 genes up-regulated in aged samples), together with TNF signaling, was more active in clusters 0 and 3 (**Fig 1e, Suppl. Fig. 3b**). These clusters were more abundant in Aged samples (**Fig. 1f**). DNA replication, E2F, and cell cycle signatures were most active in cluster 4, one of two clusters that were more abundant in Young samples (**Suppl. Fig. 3b**). Differential abundance of HSC populations between the Young and Aged cohorts was supported by cluster-independent co-varying neighborhood analysis (CNA, **Fig. 1g**). To explore this heterogeneity further, we next inferred the activity of transcription factors (TFs) using SCENIC. Despite substantial variability between datasets and individuals, combined analysis highlighted several AP-1 members as the top differentially active TFs in the aged cohort (**Fig. 1h, Suppl. Fig. 4)**, in agreement with DE results. Altogether, these observations suggest that at least part of the observed transcriptional heterogeneity in HSCs can be attributed to age, prompting us to investigate its potential sources.

To capture heterogeneity across cells that may be missed by conventional DE analysis and clustering, we used consensus non-negative matrix factorization (cNMF). This cluster-free and unbiased approach decomposes each cell’s gene expression profile into a set of underlying gene expression programs (GEPs) active across cells in the experiment (**Fig. 2a**)^20^. We ran cNMF on 22 samples with at least 100 HSCs (n=14 young, n=8 aged) to identify GEPs reproducible across datasets. We then clustered GEPs from individual samples together, which identified four representative clusters carrying at least four programs coming from at least two datasets, yielding four meta-programs (metaGEPs) (**Fig. 2b, Suppl. Fig. 5, Suppl. Table 3**). To functionally annotate metaGEPs 1-4, we assessed the enrichment of relevant gene sets in each of the meta-programs (**Fig. 2c**). metaGEP1 was enriched for the aging signature, TNF signaling, and quiescence, indicating that the activity of these processes is linked within a single program; we termed metaGEP1 the inflammatory aging program. The three remaining programs were mainly associated with lineage commitment and were annotated accordingly. metaGEP2 was enriched for a granulocyte-macrophage progenitor (GMP) signature and for cell-cycle-related pathways. metaGEPs 3-4 showed enrichment for megakaryocyte-erythrocyte progenitor (MEP) and common lymphoid progenitor (CLP) signatures, respectively (**Fig. 2c**).

Similar to our prior DE-based observations (**Fig. 1d**, **Suppl. Fig. 3b**), the activity of metaGEPs was not uniform across the population of HSCs and suggested that HSCs are composed of subpopulations of cells governed by different programs (**Suppl. Fig. 6**). To define these HSC subpopulations, we identified the predominant program in each cell of the original integrated object (**Fig. 2d**). To investigate whether the activity of metaGEPs 1-4 changes with age, we compared the fractions of cells predominated by each of the four programs in every sample between the young and aged cohorts with at least 20 HSCs (n=27 young, n=17 aged, **Fig. 2e**). Inflammatory aging metaGEP1 was more abundant in aged samples, while CLP-associated metaGEP4 showed higher activity in the young cohort. The latter agrees with lower lymphoid output with age^1,9^. Overlaying metaGEP activities with TF activities inferred by SCENIC revealed nine TFs whose activity was highly and consistently correlated with metaGEP1 usages across the datasets, including seven AP-1 complex members and two TFs implicated in maintaining quiescence, KLF4 and EGR1^21,22^ (**Fig. 2f, Suppl. Fig. 7**). These findings indicate metaGEP1 underlies age-associated differences in human HSCs and reveal molecular features of an inflammatory and quiescent HSC population that expands with age.

**Figure 2.**
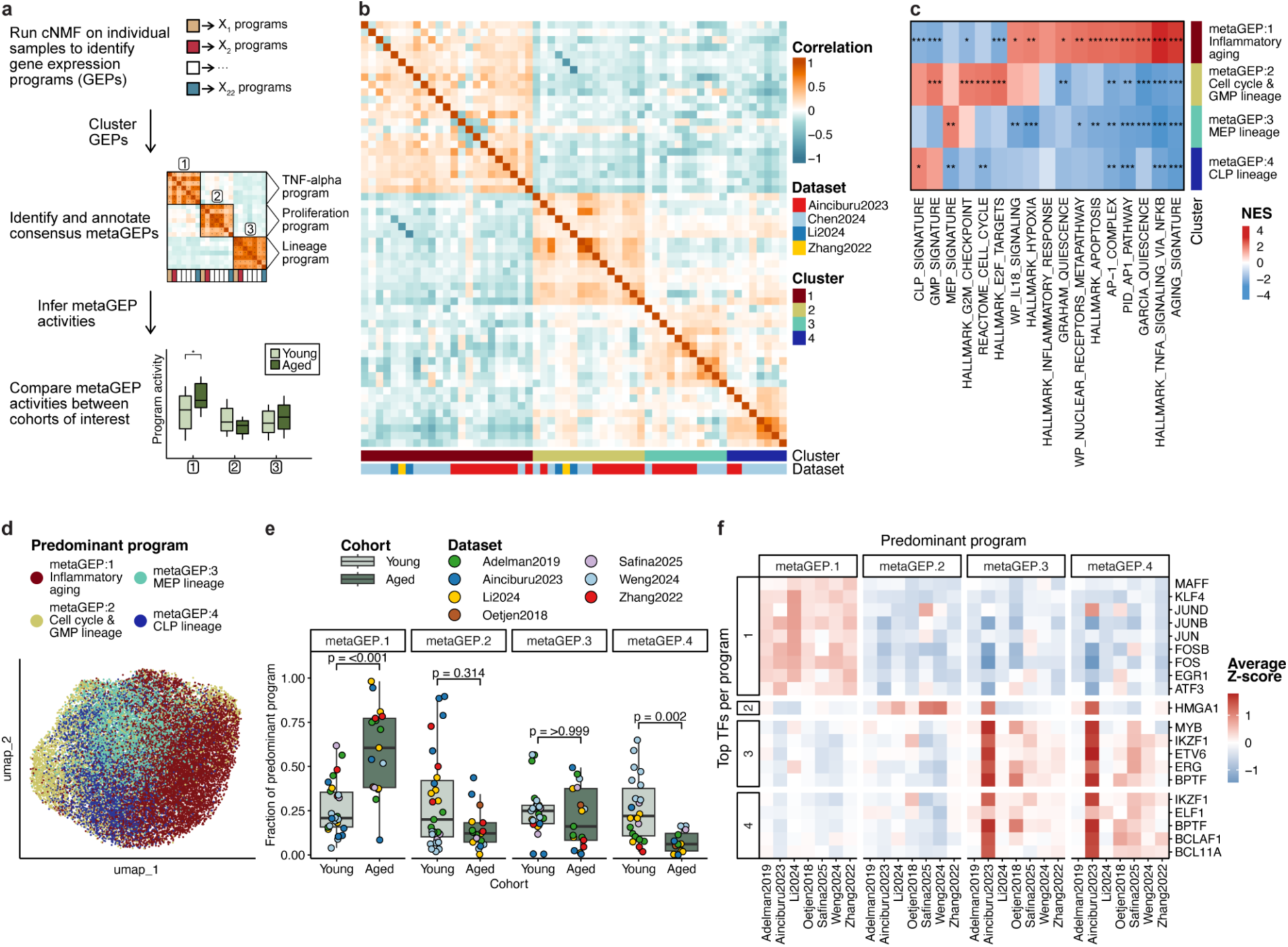
cNMF analysis recovers four gene expression meta-programs (metaGEPs) in human HSCs, including age-dependent programs. **a.** Schematic shows cNMF analysis steps. **b.** Heatmap shows Pearson correlation between Z-score vectors of individual programs comprising the four identified metaGEPs. **c.** Heatmap shows relevant pathways enriched in metaGEPs as inferred by GSEA; BH-corrected p-values: *, p<0.05, **, p<0.01, ***, p<0.001. **d.** UMAP shows all integrated HSCs, colored by the predominantly active metaGEP. **e.** Box plot shows comparison of predominant program abundances between the Young (n=27) and Aged (n=17) cohorts. Symbols indicate cluster fractions per sample; p-values of two-sided Wilcoxon test with Bonferroni correction are shown. **f.** Heatmap shows activity of metaGEP-specific regulons, identified by SCENIC; averaged Z-scores of AUC values are shown. GEP: gene expression program, GMP: granulocyte-macrophage progenitor, MEP: megakaryocyte-erythrocyte progenitor, CLP: common lymphoid progenitor.

In this study, we aggregated data from seven single-cell and single-nucleus studies and consistently annotated 28,989 HSCs across individuals, including 27 young and 17 aged samples with at least 20 HSCs, and identified four meta-programs that drive transcriptional heterogeneity through variable activity across HSCs. The largest meta-program we recovered, metaGEP1, was enriched for TNF signaling and quiescence, and marked a subpopulation of HSCs that expanded with age. These findings indicate that HSC aging is not uniform but is instead shaped by shifting program activities.

Inflammation has been linked to HSC activation and exhaustion^23–25^. In contrast, our data link inflammatory signaling and quiescence within a single program associated with aging, more reminiscent of chronic inflammation which can induce quiescence^26,27^. With the emerging concept that acute inflammatory stimuli induce trained immunity in HSCs^13,28,29^, an outstanding question is to what extent metaGEP1-dominated HSCs generate pro-inflammatory immune cells that contribute to aging.

Members of the AP-1 complex were among the top-enriched transcription factors in both DE and cNMF analysis. While the activity of immediate early response (IER) genes, including members of AP-1, can be prone to technical variation due to sample processing^30^, our finding that AP-1 genes are consistently upregulated across several datasets implies biological relevance. Along with other genes highlighted in our analysis, including anti-inflammatory ZFP36 and NFKBIA^31,32^, perturbation of these factors may modulate HSC aging.

The cluster-independent framework we used to discover the biology of aging HSCs can be easily adapted to larger datasets. Indeed, there were only 22 samples with >100 HSCs from which to discover GEPs, and larger samples tended to resolve more programs successfully (**Suppl. Fig. 8**). Analyzing more samples with larger cell counts will improve the numerical stability of NMF solutions and may allow for more granular resolution of HSC programs. Finally, since inflammatory pathways in HSCs are affected by clonal hematopoiesis^14^, future work should assess how genetic changes interact with the programs identified here.

## Conclusions

Our study provides a consensus, single-cell resolution map of human HSCs by integrating seven prior datasets. Among the most robust changes in aging HSCs are upregulation of pathways associated with inflammatory signaling (TNF, IFN-g), stress-activated MAPK signaling (AP-1), and quiescence. We use a cluster-free approach to uncover activity programs that vary in intensity across individual HSCs. The strongest program shaping HSC heterogeneity is associated with inflammation and quiescence, linking these properties together in individual cells. An inflammatory HSC subset dominated by this program expands with age. These findings open new directions for testing whether modulating inflammatory HSCs can rejuvenate blood production and help maintain healthy hematopoiesis with age.

## Declarations

### Ethics approval and consent to participate

Sternum bone marrow samples from the Safina dataset were collected under an excess sample banking and sequencing protocol approved by the Mass General Brigham Institutional Review Board (IRB), which covered all study procedures. Additional datasets were obtained from public repositories.

### Consent for publication

Not applicable.

### Availability of data and materials

Data are available at https://figshare.com/projects/Aging_HSCs_2025/235781 (under embargo until publication). Annotated code to reproduce all analyses is available at https://github.com/noranekonobokkusu/Aging_HSCs_2025 and will be made available upon publication. Sequencing data generated in this study were deposited to GEO under accession number GSE302126 and will be made available upon publication.

### Authors’ contributions

Conceptualization and study design: K.R.S., P.v.G. Supervision: P.v.G. Sample and data collection: K.R.S., J.D.G., C.W., A.K., V.G.S. Analysis and visualization: K.R.S. Results interpretation: K.R.S., D.A.K., M.C., J.T., V.G.S., P.v.G. Manuscript draft: K.R.S., P.v.G.. Manuscript review and editing: K.R.S., D.A.K., M.C., J.D.G., C.W., S.D., S.R., A.K., J.T., V.G.S., P.v.G. All authors read and approved the final manuscript.

## Acknowledgments

We thank members of the Van Galen Laboratory for helpful discussions. We thank Nathan Salomonis, Christopher Hourigan, and Hojun Li for instructions on data usage.

## Methods

### Single-cell sequencing

We generated one of the seven single-cell RNA-seq datasets ourselves (others were publicly available). Bone marrow cells from nine donors were collected from the iliac crest of patients undergoing cardiac surgery under an excess sample banking and sequencing protocol that covers all study procedures and was approved by the Institutional Review Board (IRB) of Mass General Brigham. Donors were confirmed negative for common CHIP mutations using targeted sequencing. Mononuclear cells were isolated using Ficoll or lymphoprep and cryopreserved in liquid nitrogen storage. Cells were thawed using standard procedures, and viable (DAPI negative) cells were sorted on a Sony SH800 flow cytometer. Next, 10,000-15,000 cells were loaded onto a 10x Genomics chip. Further processing was done using the recommended procedures for the 10x Genomics 3’ v3.0, v3.1, or v4 chemistry. Libraries were sequenced on the NovaSeq SP 100 cycle with the following parameters (Read 1: 28 + Read 2: 75 + Index 1 (i7): 10 + Index 2 (i7): 10). Count matrices were generated using CellRanger v.7.1.0 with default settings and GRCh38 as the reference genome.

### Dataset preparation and annotation

We compiled six publicly available human single-cell datasets and added nine samples from our lab. The complete dataset contained bone marrow cells from 98 individuals (9 prenatal, 2 cord blood, 4 infant, 7 child, 31 young, 15 middle-aged, and 30 aged)^4,11,33–36^. Gene expression matrices were available for each of the datasets, except for the Adelman and some samples in the Weng datasets. For the Adelman dataset, we mapped sequencing data onto hg38 using STAR^37^ with default settings and quantified gene counts using featureCounts from Rsubread package^38^ to produce a gene expression matrix. For the Weng dataset, we prepared the input object as described in Github repository petervangalen/ReDeeM_2024. For each of the seven datasets, we loaded the count matrix into Seurat^39^, keeping genes captured in at least five cells and cells with at least 100 genes captured. We then filtered the initial matrix based on nCount, nGene, mitochondrial genes content, and doublet scores (as determined by scrubletR^40^) for each sample in the dataset, removing differentiated cell populations where applicable. The accession codes and dataset-specific filtering parameters are available in **Supplementary Table 1**. We then subsetted each of the Seurat objects for genes present in all of the datasets (n=11,612). To obtain concordant cell type annotations among the datasets, we mapped each of the datasets onto the BoneMarrowMap atlas and transferred cell type labels^41^. For subsequent analyses, we only retained cells annotated as HSCs. 30,070 cells were annotated as HSCs; of those, 1,081 cells were mapped outside the reference HSC cluster and were excluded from the analysis, leaving us with 28,989 HSCs.

To cross-validate BoneMarrowMap-based HSC assignment, we integrated seven HSPC datasets with scVI^42^ (v1.3.0, n_layers=4, n_latent=30, max_epochs=60), specifying sample name as a batch key and dataset name and single-cell vs. single-nucleus data type as categorical covariates (batch_key=’Sample’, categorical_covariate_keys=[’Dataset’, ’data_type’]), uploaded the integrated object to Seurat, computed neighborhood graph on scVI latent variables, identified Louvain clusters (resolution=1.5) and computed UMAP. We scored each cell by three published HSC signatures^43–45^, computed average signature scores per cluster (aggregating across cells and three public signatures), and selected four clusters with the highest average HSC score as HSCs. This yielded 44,187 cells and included 87% of HSCs identified by BoneMarrowMap, implying the two annotation approaches are concordant, with BoneMarrowMap being more specific.

### Differential expression analysis

To identify gene expression changes associated with age, we pseudobulked HSC samples per sample and ran DESeq2^46^ using dataset name and sample sex as covariates. Race and ethnicity were not reported in publicly available datasets and therefore not included as covariates. To ensure the robustness of the analysis, we only used samples with at least 20 cells and datasets with at least two such samples in both Young and Aged cohorts (n=25 young and n=15 aged samples across five datasets). We obtained results for the ‘Cohort_Aged_vs_Young’ coefficient and shrunken log fold changes using apeglm^47^. To define the aging signature, we used genes with logFC>1 and p.adj<0.05, yielding 68 genes.

### Gene set enrichment analysis

We used the fgsea package^48^ to conduct gene set enrichment analysis (GSEA) and characterize ranked gene lists produced in this study. We used two collections of signatures from MSigDB^49–51^, HALLMARK and C2:CP. We obtained a quiescence signature from García-Prat et al. by selecting genes with logFC < -2 and FDR<0.00001 from the DE results of Table S1 therein^52^. To define MEP, GMP, and CLP lineage signatures, we identified markers of MEP, Early GMP, and CLP populations of the BoneMarrowMap-annotated object, respectively (using Seurat’s FindMarkers function, providing HSC, MPP-MkEry and MPP-MyLy as ident.2). For each lineage, we selected top-50 genes with p.adj<1E-10 and logFC>1, and excluded genes overlapping between the three 50-gene sets. To annotate cNMF metaGEPs, we also included the aging signature (68 genes) identified in the DE analysis of this study. To characterize the recovery of the aging signature compared to a random signature, we generated 20 random signatures with an expression pattern similar to the aging signature and included them in GSEA (as detailed below).

### Variable genes identification

To identify variable genes shared across the seven datasets, we first excluded ribosomal genes (n=94 genes starting with RPS or RPL), then identified top 3,000 variable features within each dataset using FindVariableFeatures() in Seurat, and finally, selected the genes identified as variable in at least 4 datasets out of 7 (n=1,554). We then subsetted the expression count matrix for these genes and used it as an input for scVI and cNMF.

### Dataset integration

We integrated datasets using scVI (v1.3.0, n_layers=3, n_latent=30), specifying sample name as a batch key and dataset name as a categorical covariate (batch_key=’Sample’, categorical_covariate_keys=[’Dataset’]), uploaded the integrated object to Seurat, computed neighborhood graph on scVI latent variables, identified Louvain clusters (resolution=0.5) and computed UMAP. Louvain clusters and UMAP coordinates were transferred to the original object with the entire gene set (11,612 genes), to score cells for gene sets of interest using AddModuleScore() (**Fig. 1d**, **Suppl. Fig. 3a**).

Consensus non-negative matrix factorization for inference of gene expression programs We used consensus non-negative matrix factorization (cNMF)^20^ to identify gene expression programs (GEPs) across datasets. To mitigate the effect of sample-driven batch effects, we identified GEPs at the level of individual samples and clustered the discovered programs to define meta-programs (metaGEPs) shared across multiple samples and datasets, similar to the approach taken by Gavish et al.^53^ We used young or aged samples with at least 100 cells, resulting in 8 Ainciburu, 6 Weng, 7 Li, and 1 Zhang sample (total n=22). As cNMF is sensitive to cells with low gene counts and low number of genes, we additionally preprocessed each of the samples as follows. We preserved cells with at least 50 captured genes and the total number of transcripts within the sample-specific limits (see below), and genes expressed in at least 10 cells. Sample-specific total count limits were defined as median+/-2.5 mean absolute deviation for each sample except sample BM3 in Zhang dataset. Data from Zhang et al. has an order of magnitude higher expression counts than other datasets with several cells having much higher counts than the median; these outlier cells drive individual programs in cNMF results, dominating the observed variation in expression data. To address this, we used a fixed interval for total counts of [100-10,000] for Zhang sample BM3.

After preprocessing, we ran cNMF on each sample, with 500 iterations of factorization and a k (the number of inferred GEPs) ranging from 3 to 10. For each sample, a single final value of k was selected based on the stability vs. error plot and visual inspection of 500 clustered factorization results. We assessed distance thresholds of 0.05, 0.1 and 0.15; distance threshold of 0.1 produced the most visually stable clusters and was used for all the samples. Among the smaller samples, individual cells sometimes drive some of the identified programs, similar to sample BM3 in Zhang dataset. To address this, for each program, we computed the ratio between 100% and 75% quantiles of usage values, and excluded programs with a ratio of more than 10 (which reflects that the maximum usage value is much larger than most usage values). After this filtering, 97 GEPs remained.

### Identification and analysis of metaGEPs

We clustered Z-score spectra of 97 GEPs using the iterative clustering algorithm implemented in starCAT^54^ with a modification allowing for more than one program per sample in a cluster (corr_thresh=0.1, pct_thresh=0.1). We defined meta-programs (metaGEPs) as clusters carrying at least four programs coming from at least two datasets. Z-score spectra within each metaGEP were averaged and annotated with GSEA. The variance-normalized TPM spectra within each metaGEP were averaged to produce the meta-program expression matrix (4 metaGEP x 11,612 genes) which we used to infer metaGEP usages in each of the datasets with starCAT^54^. To compare usages between the young and aged cohorts, we defined a single predominant metaGEP per cell, computed the fractions of each predominant metaGEP in every sample with at least 20 cells (n=27 young and n=17 aged samples), and compared the fractions between the two age cohorts.

The aging signature was the most consistently recovered gene set, as it showed the most frequent significant enrichment by GSEA on GEPs. To test whether this high recovery rate is specific to the aging signature rather than its gene expression pattern, we constructed 20 random gene sets and assessed their recovery by GSEA. To construct a random gene set, we computed average normalized expression for 11,612 genes, split the genes into 50 equidistant expression bins, and randomly sampled 68 genes from the bins containing aging signature genes, to mimic the expression pattern of the aging signature. To assess the recoverability of gene sets by GSEA, we revisited our cNMF results, selected all visually stable cNMF runs (k values with distance thresholds producing the most stable clusters), and annotated them with GSEA using 20 random gene sets, the aging, TNF signaling, quiescence signature, and the three lineage signatures, applying a Bonferroni correction (which is more stringent than the default Benjamini-Hochberg procedure). For each cNMF run within each sample, we assessed whether it contains a program significantly positively enriched for any of the tested gene sets, and computed the fraction of cNMF runs within each sample that recovered each of the gene sets (**Suppl. Fig. 8a**). The aging signature was indeed the most recoverable; lineage signatures were less recoverable, but invariably still more recoverable than the random gene sets. We also assessed the recovery rate of a combination of gene sets (Aging + TNF + Quiescence, and Aging + TNF, **Suppl. Fig. 8b**), which also showed higher recovery than random gene sets.

Samples with more HSCs had higher recovery rates than samples with fewer HSCs. This demonstrates that the genesets we used to annotate metaGEPs are robust compared to random gene sets, supporting biological significance, and that using additional large (CD34+ enriched) samples may help resolve and annotate gene expression programs better.

### SCENIC analysis

We used pySCENIC to infer the activity of transcription factors (TFs)^55^. We followed the standard pipeline with default parameters, except using ‘--auc_threshold 0.01’ for the ‘ctx’ command. We inferred TF activities (represented as area under the curve values, AUC) within each dataset separately and then retained 108 TFs identified in all seven datasets. To identify age-specific TFs, we first subsetted young and aged samples, converted age cohort to a binary variable (with 1 being Aged), and computed point-biserial correlation between the binary age variable and TF activities within each dataset. We then assigned ranks to TFs within each dataset, averaged both ranks and correlation values, and selected TFs with mean rank <=10 and mean correlation >=0.15 as TFs associated with age, yielding five TFs (**Fig. 1h**). We repeated the same procedure to infer TFs associated with each of the metaGEPs, retaining samples from all age cohorts and computing Pearson correlation between TF activities and metaGEP usages, instead of point-biserial correlation (**Fig. 2f, Suppl. Fig. 7**). For visualization purposes, AUC values were converted into Z-scores. We limited the Z-score values to the 0.1 and 99.9 percentiles when visualizing TF activities in individual cells (**Suppl. Fig. 4**) to avoid distorting the color scale with outliers.

### Co-varying Neighborhood Analysis

To test whether the age cohort is associated with certain neighborhoods of cells, we conducted association testing using co-varying neighborhood analysis (CNA)^56^. We first subsetted the integrated object for young and aged samples, encoded age as a binary variable (with 1 being Aged), computed a neighborhood graph using scVI latent variables, and ran CNA, correcting for the dataset name as a batch variable. To visualize significant associations in the UMAP, we colored neighborhoods with FDRs>0.1 grey.

## Supplementary Figures

**Supplementary Figure 1.**
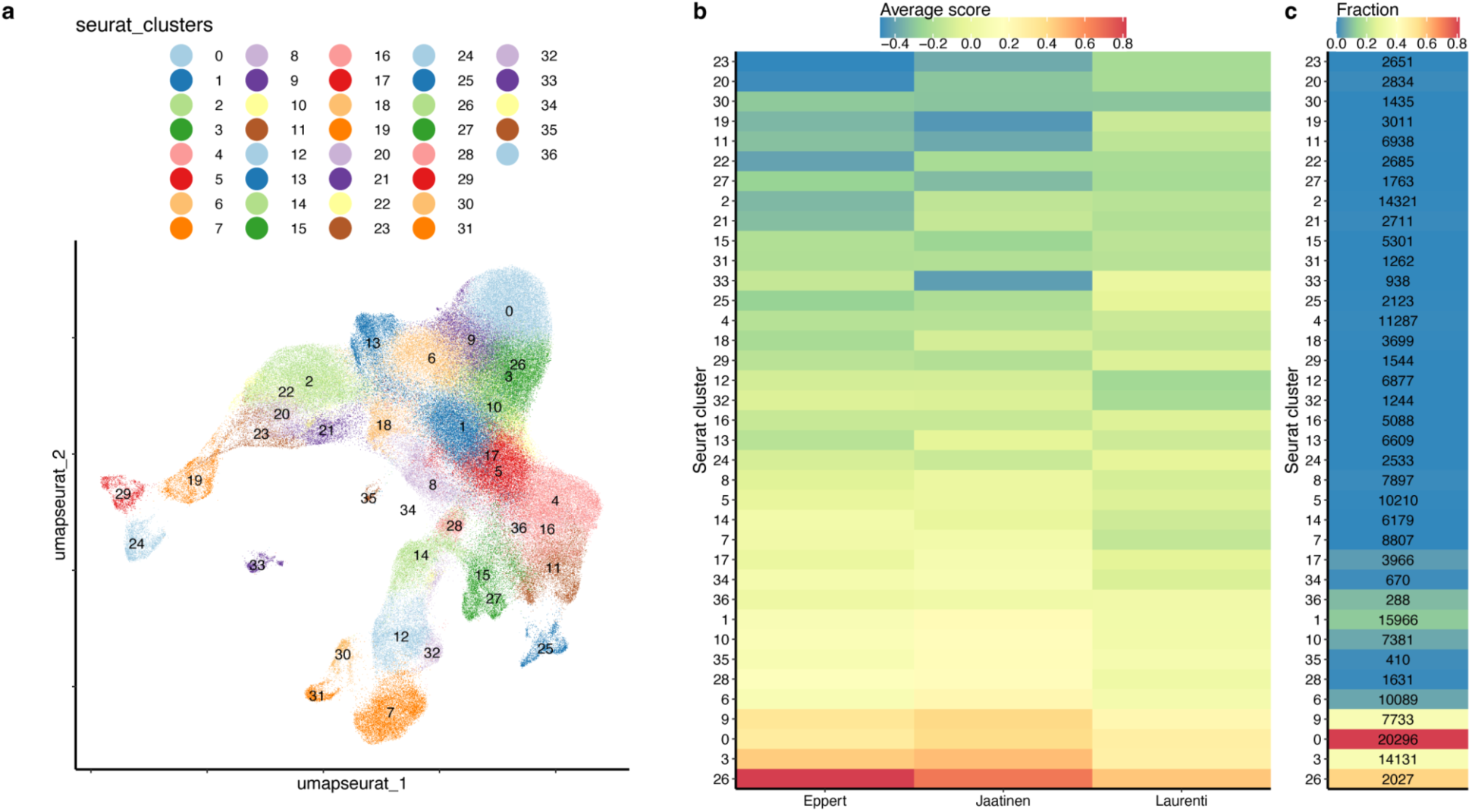
Orthogonal HSC annotation agrees with BoneMarrowMap-produced HSC annotation. **a.** UMAP shows Louvain clusters identified on the integrated HSPC object. **b.** Heatmap shows the average module scores of three HSC-related public signatures across the Louvain clusters; clusters are sorted by the average score across the three signatures. **c.** Heatmap shows the fraction of cells annotated as HSCs by BMM per cluster; text shows the total number of cells per cluster. Clusters 0, 9, 3, and 26 sum up to 44,187 cells which includes 87% of the 28,989 BoneMarrowMap-annotated HSCs.

**Supplementary Figure 2.**
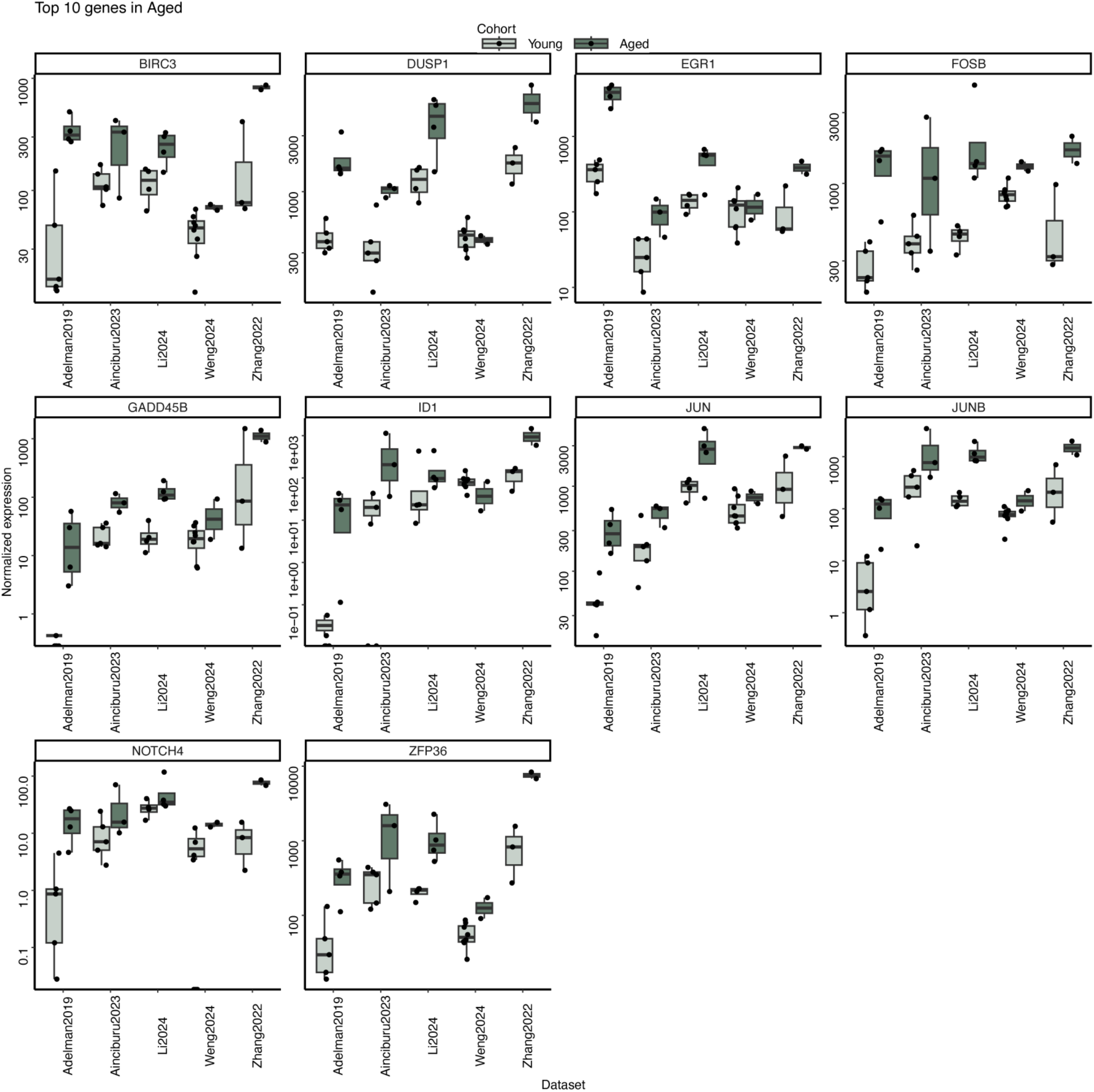
Differential gene expression analysis reveals age-associated expression changes. Box plots show normalized expression of top-10 genes most significantly up-regulated in the Aged cohort across five datasets used for DE analysis. Each symbol represents an individual donor sample.

**Supplementary Figure 3.**
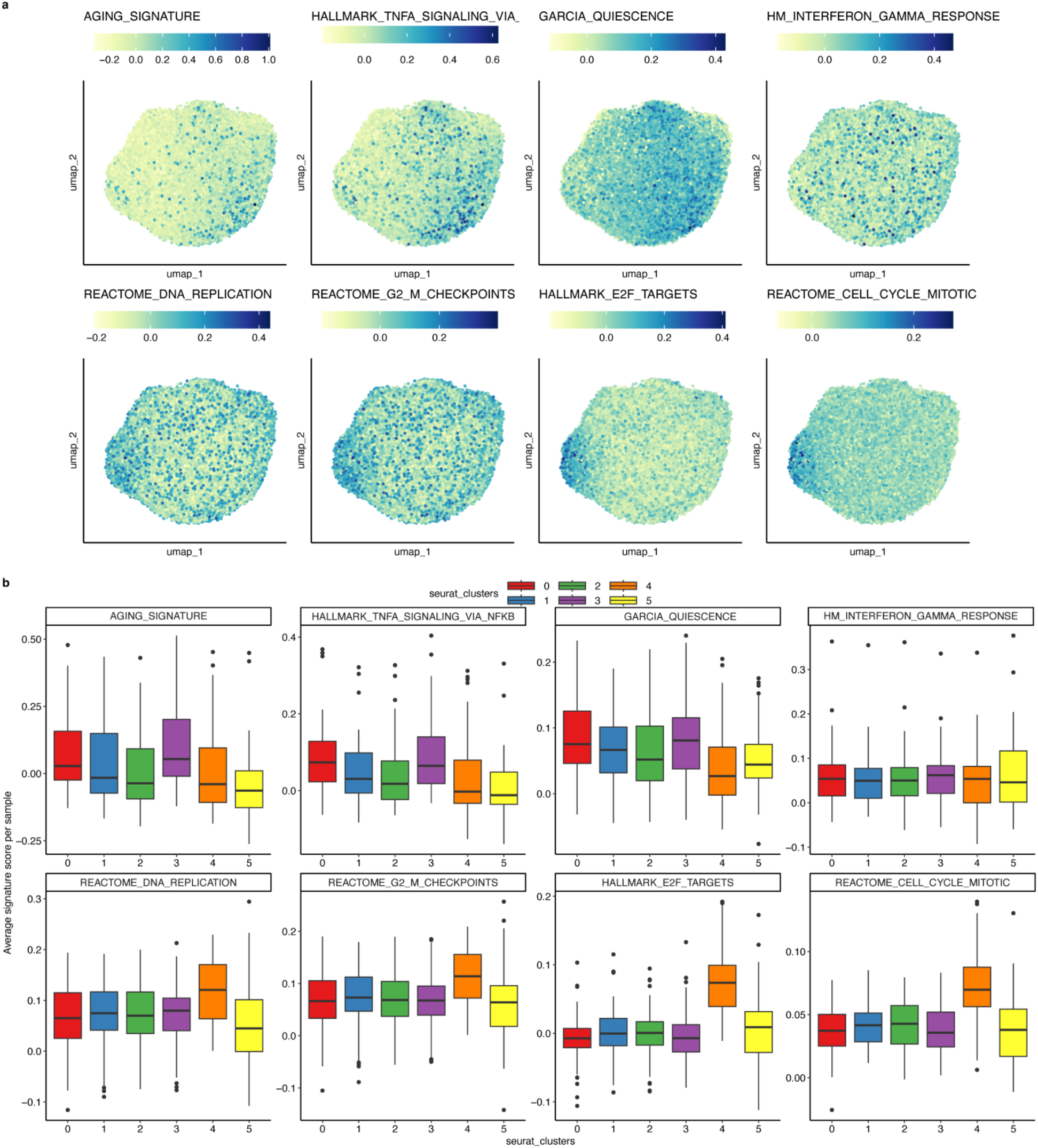
The activity of representative gene sets up-regulated in Aged (top row) and Young (bottom row) samples is not uniform across the population of HSCs. **a.** UMAPs colored by signature scores. **b.** Average signature score per sample across Seurat clusters. Only samples with at least 20 cells are used.

**Supplementary Figure 4.**
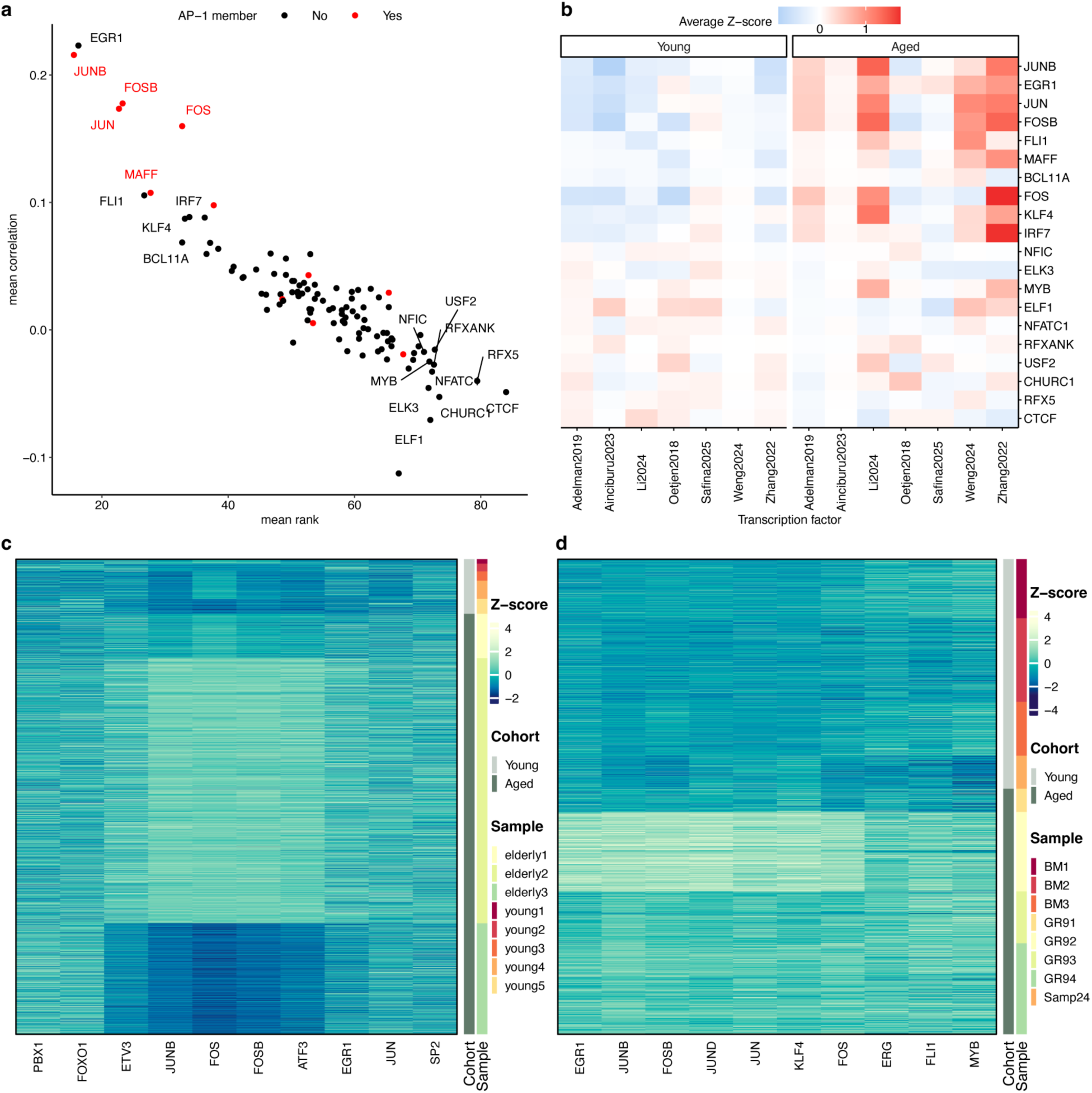
Combined analysis of age-associated transcription factor activities. **a.** Scatter plot shows the mean ranks (x) vs. mean correlation (y) with age status of 108 TFs recovered in SCENIC analysis. Top-10 and bottom-10 TFs are labeled, AP-1 members are marked red. **b.** Heatmap shows the average Z-score of TFs labeled in (a) across all datasets. **c, d.** Heatmaps show the activity of top-10 age-associated TFs (columns) in cells (rows) from two example datasets: Ainciburu (c) and Li (d). Significant variability between samples can be observed, including samples within the same age group.

**Supplementary Figure 5.**
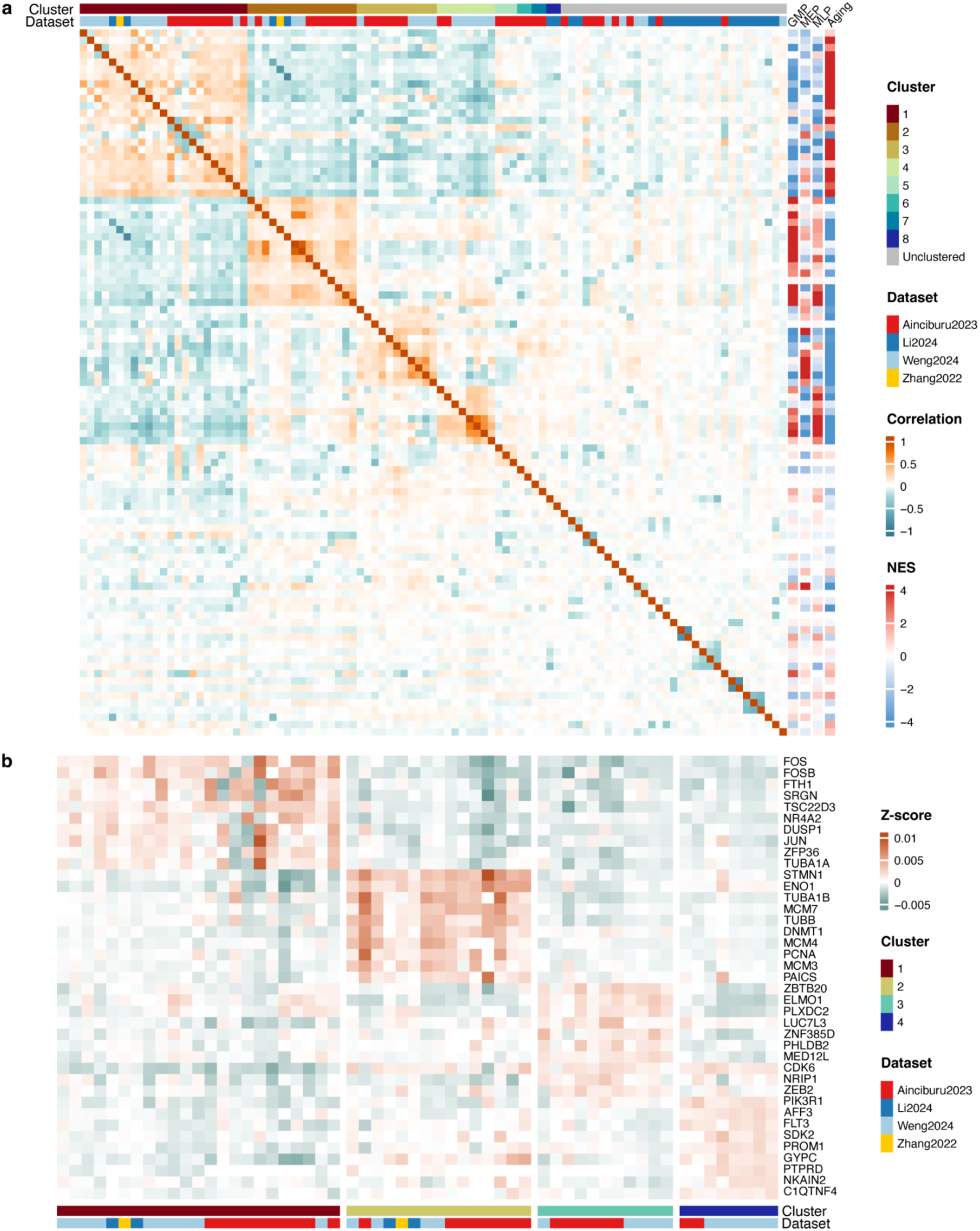
cNMF analysis infers four metaGEPs. **a.** Heatmap shows clustering of all 97 programs discovered across 22 samples. This is an extended version of Fig. 2b. The side bars show NES for characteristic signatures, indicating consistent enrichment of signatures across multiple programs within clusters. **b.** Heatmap shows Z-scores for top-10 genes associated with each cluster (metaGEP), highlighting consistent enrichment across most programs within each cluster.

**Supplementary Figure 6.**
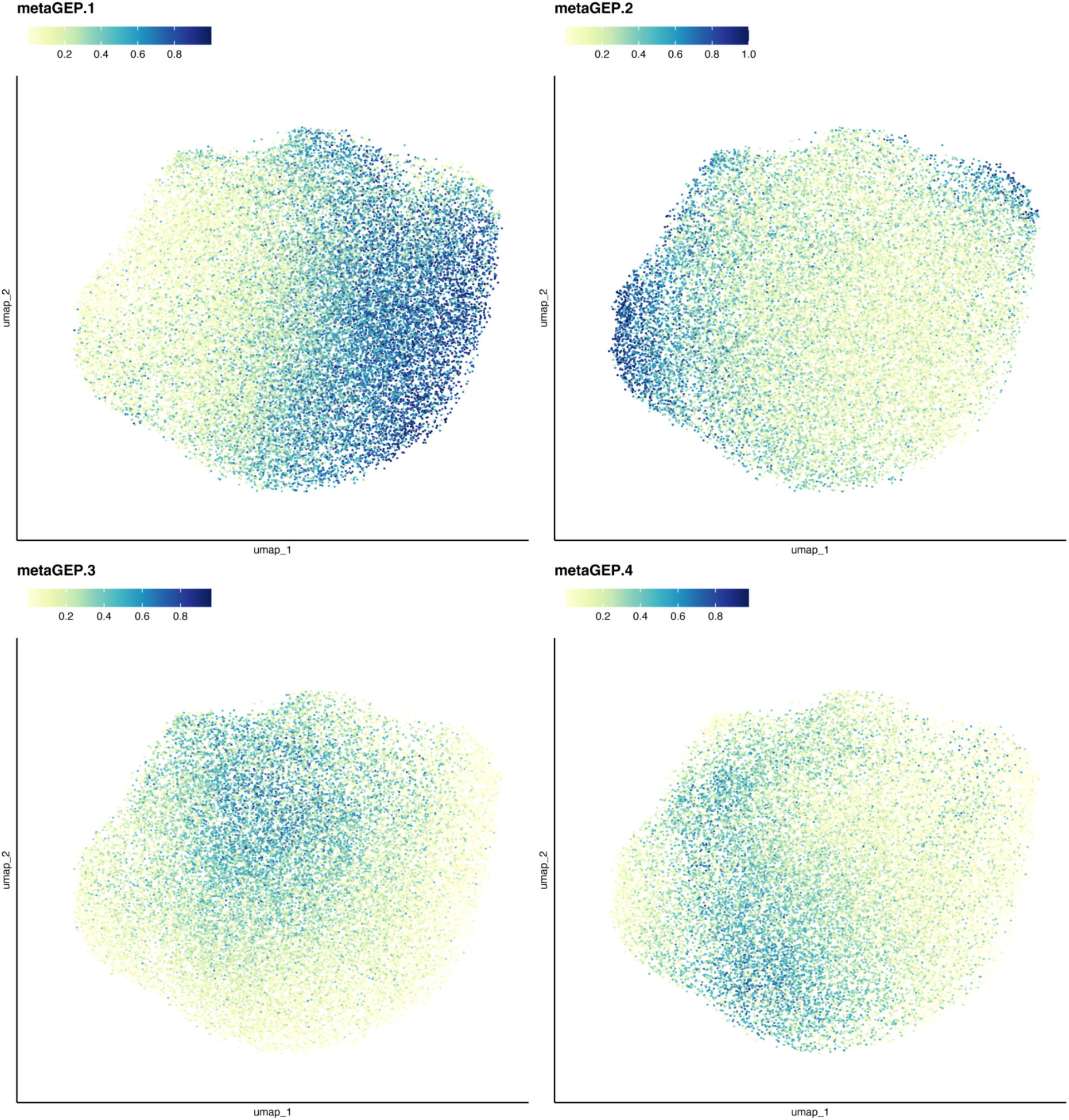
Activities of metaGEPs are not uniform across HSCs. UMAPs are colored by usages of metaGEPs 1-4, normalized to a total of one by starCAT.

**Supplementary Figure 7.**
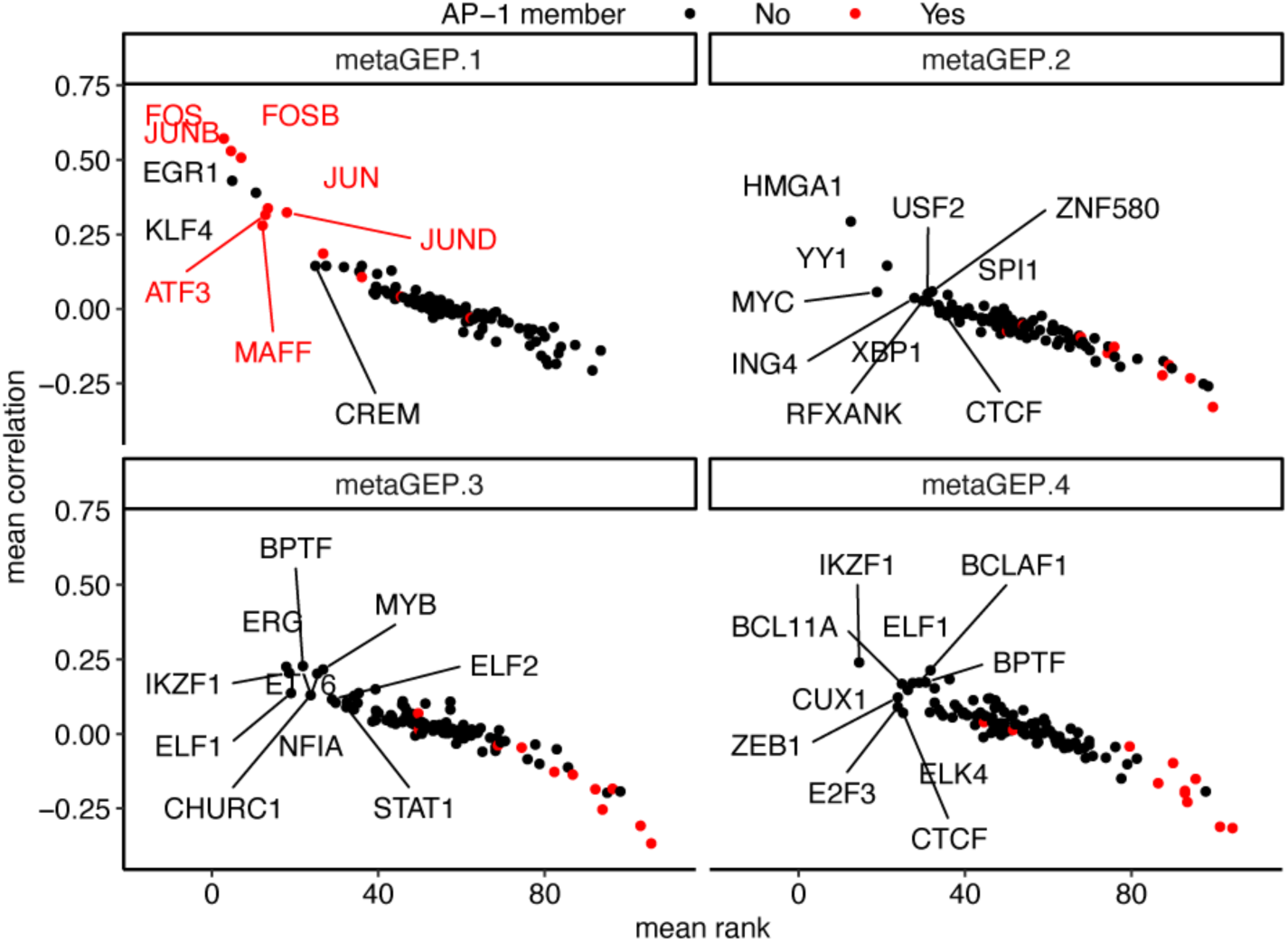
metaGEPs are characterized by specific TF activities. Scatter plots show TFs (symbols) by their average rank across seven datasets (x) vs. their average correlation of AUC scores and metaGEP usages (y). Top-10 TFs per metaGEP are labeled; AP-1 members are shown in red. TFs with mean rank <=10 and mean correlation >=0.15 were selected for visualization in Figure 2f. Compared to metaGEP1, correlation of TF activities with metaGEPs 2-4 was weaker, revealing overlapping TFs, particularly between metaGEPs 3-4. Thus, in contrast to the easily recoverable inflammatory aging program, metaGEP1, lineage priming programs may require more data to be reliably inferred and disentangled at the level of TF activity.

**Supplementary Figure 8.**
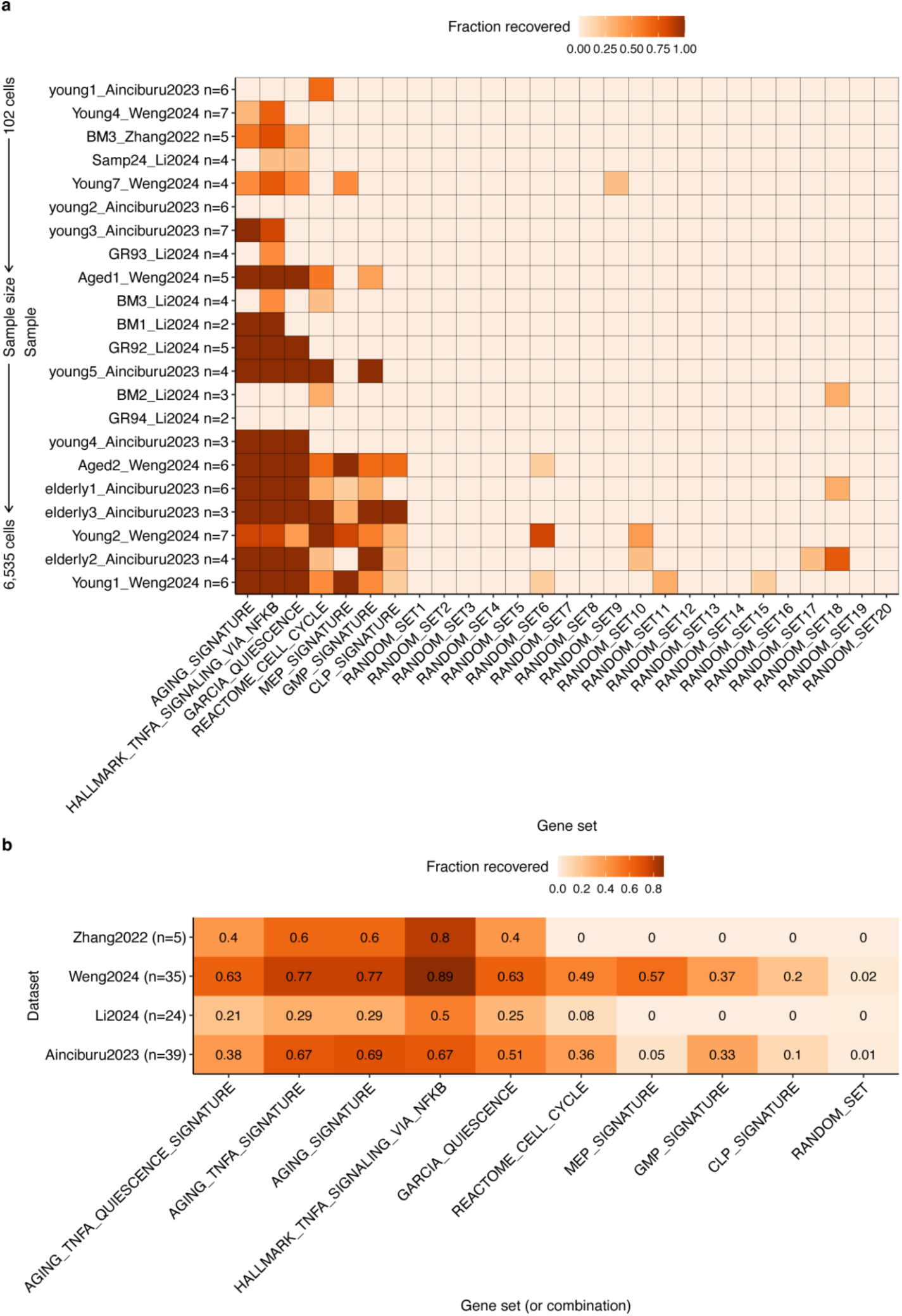
Functionally relevant gene sets and their combinations are enriched in cNMF programs, compared to 20 random gene sets. **a.** Fraction of stable cNMF runs that recovered a positively enriched gene set, across 22 samples used for cNMF; samples are ordered by the number of HSCs, ***n*** specifies the number of visually stable cNMF runs per sample (for example, ‘Young1_Weng2024 n=6’ means that for sample Young1 from dataset Weng2024, there were six stable cNMF runs among k=3..10 tested). **b.** Recovery rate of gene sets/their combinations, aggregated across datasets; ***n*** specifies the total number of stable cNMF runs per dataset, numbers in tiles show fractions; for the random set, the average fraction across 20 random sets is shown.

